# Segmented K-Space Blipped-Controlled Aliasing in Parallel Imaging (Skipped-CAIPI) for High Spatiotemporal Resolution Echo Planar Imaging

**DOI:** 10.1101/2020.06.08.140699

**Authors:** Rüdiger Stirnberg, Tony Stöcker

## Abstract

**Purpose:** A segmented k-space blipped-CAIPI (skipped-CAIPI) sampling strategy for echo planar imaging (EPI) is proposed, which allows for a flexible choice of EPI factor and phase encode bandwidth independent of the controlled aliasing (CAIPIRINHA) pattern.

**Theory and Methods:** With previously proposed approaches, exactly two EPI trajectories were possible given a specific CAIPIRINHA pattern: either with slice gradient blips (blipped-CAIPI), or following a shot-selective approach (higher resolution). Recently, interleaved multi-shot segmentation along shot-selective CAIPI trajectories has been applied for high-resolution anatomical imaging. For more flexibility and a broader range of applications, we propose segmentation along any blipped-CAIPI trajectory. Thus, all EPI factors and phase encode bandwidths available with traditional segmented EPI can be combined with controlled aliasing.

**Results:** Temporal signal-to-noise ratios of moderate-to-high-resolution time series acquisitions at varying undersampling factors demonstrate beneficial sampling alternatives to blipped-CAIPI or shot-selective CAIPI. Rapid high-resolution scans furthermore demonstrate SNR-efficient and motion-robust structural imaging with almost arbitrary EPI factor and minimal noise penalty.

**Conclusions:** Skipped-CAIPI sampling increases protocol flexibility for high spatiotemporal resolution EPI. In terms of signal-to-noise ratio and efficiency, high-resolution functional or structural scans benefit vastly from a free choice of the CAIPIRINHA pattern. Even at moderate resolutions, the independence of sampling pattern, echo time and image matrix size is valuable for optimized functional protocol design. Although demonstrated with 3D-EPI, skipped-CAIPI is also applicable with simultaneous multislice EPI.

## Introduction

An increasing number of structural imaging (1–6), quantitative imaging (7–10) or high spatiotemporal resolution functional imaging (11–21) applications have been approached by dedicated echo planar imaging (EPI) (22) implementations. Many of those aim at higher spatial resolution than usual, e.g. in functional MRI (fMRI). It is generally appealing to combine the inherent signal-to-noise ratio (SNR) efficiency of EPI (23,24) with controlled aliasing in parallel imaging (CAIPIRINHA, short: CAIPI (25)) to minimize the geometry-dependent parallel imaging noise penalty (g-factor (26)). However, previous sampling approaches limit each CAIPI pattern to only two possible EPI k-space trajectories and corresponding EPI factors: single-shot blipped-CAIPI (27) and multi-shot shot-selective CAIPI (18,28,29). Thus, higher-priority parameters affecting the EPI factor as well (e.g. base resolution and echo time) often enforce a suboptimal CAIPI pattern instead of a noise-optimal one. Recently, shot-selective CAIPI has been combined with interleaved multi-shot segmentation fostering rapid T_1_-weighted anatomical imaging (6). Still, this approach is limited to only a fraction of traditional segmentation options without controlled aliasing. We demonstrate that introducing a segmentation factor into blipped-CAIPI overcomes these limitations and proves useful for a wide range of high spatiotemporal resolution applications.

## Theory

### Segmented EPI

Interleaved multi-shot segmentation has been proposed very early to achieve higher resolutions with reduced off-resonance artifacts (e.g. geometric distortions, chemical shift) and T_2_*-related artifacts (“T_2_* blur”), shorter echo times (TE) and repetition times per shot (TR) (30). Parallel imaging along the blipped phase encode direction (*y*, w.l.o.g.) also helps with this regard: combined with *S*-fold segmentation, the phase encoding *y*-blip becomes *S · R_y_*, compared to only *R_y_* (undersampling factor along *y*) without segmentation. The *y*-blip directly controls the EPI factor, *EF* = ┌*N_y_*/(*S* · *R_y_*┐, and the phase encode bandwidth, *BW_y_* = *S* · *R_y_*/*ESP*, where *N_y_* and *ESP* denote the phase encode matrix size and the echo spacing, respectively. Echo time shifting can be introduced to smooth out signal magnitude and phase jumps between adjacent odd-/even-echo k-space sections (31,32). The segmentation technique has been combined with 3D-EPI and with parallel imaging along two directions (Fig. 1 A), e.g. for T_2_*/susceptibility-weighted high-resolution imaging (1,2) or fMRI (13–15,17,19,33), but these implementations lacked controlled aliasing.

**Figure 1:**
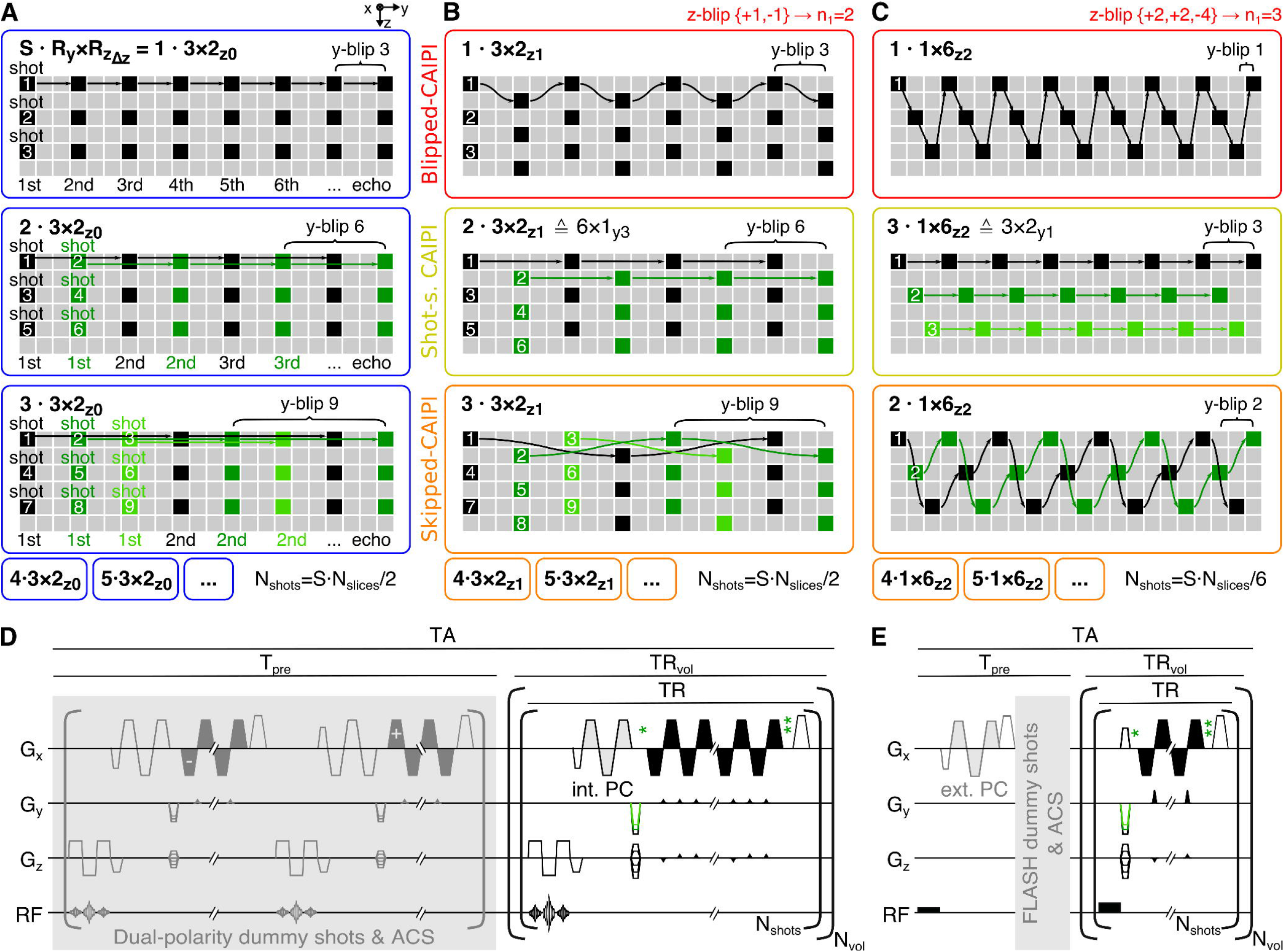
Schematic depiction of traditional (segmented) 3D-EPI k-space sampling with 3 × 2 undersampling (A) and derived CAIPI trajectories for the corresponding 3 × 2_z1_ CAIPI pattern (B) and for an alternative 1 × 6_z2_ CAIPI pattern (C). Row 1 and 2: the blipped-CAIPI and the shot-selective CAIPI trajectory for the respective pattern. Row 3: one skipped-CAIPI trajectory example. D and E: schematic diagram of slab-selective (oblique-axial time series) and whole-head (sagittal high-resolution) sequence variants used here. Pre-(*)/post-(**)-EPI delay times increase/decrease with the first primary phase encode value per shot for echo time shifting (see Supporting Information Figure S8 for visualization). N_shots_=S·N_slices_/R_z_: number of shots per volume measurement. N_vol_: number of volume measurements. N_slices_: number of slices. PC: phase correction scan. ACS: autocalibration scan.

### Blipped-CAIPI

Simultaneous multislice (SMS) EPI (27) or 3D-EPI using blipped-CAIPI sampling (16,20,34–36) can employ stronger undersampling owing to a reduced g-factor. Positive and negative gradient blips along the slice or secondary phase encode axis (*z*, w.l.o.g.) are introduced between echoes along the EPI train, resulting in a certain CAIPI sampling pattern

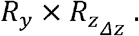

Here, *R_z_* and *Δz* denote the undersampling (multiband) factor and the CAIPI shift along *z*, respectively. The same pattern can also be expressed with a CAIPI shift along *y*. The *z*-notation (37) is adopted here, with −└*R_z_*/2┘ ≤ *Δz* ≤ └*R_z_*/2┘, Generally, selecting an adequate CAIPI pattern minimizes the g-factor in the fundamental relation (26):

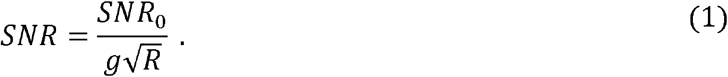

*SNR*_0_ is the volumetric SNR without undersampling and

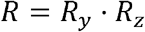

is the total undersampling factor. For SMS-EPI, 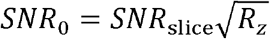 is often inserted into Eq. 1, where *SNR*_slice_ is the single-slice SNR without undersampling. Either way, each CAIPI pattern corresponds to one unique blipped-CAIPI trajectory. In *z*-notation, the *y*-blip is exactly *R_y_* (Fig. 1, row 1).

### Shot-selective CAIPI

At high spatiotemporal resolutions it has been proposed to acquire only those CAIPI samples in a single shot, that correspond to one secondary phase encode value. In the next shot, all samples that correspond to the next secondary phase encode value are acquired, etc. (28,29). Such a shot-selective CAIPI approach has recently been applied with SMS-EPI (5) and 3D-EPI (18) to achieve greater *y*-blips. Each CAIPI pattern corresponds to one unique shot-selective trajectory without *z*-blips. The *y*-blip is exactly *R_z_* times larger than that of the blipped-CAIPI trajectory for the same CAIPI pattern (Fig. 1 B, row 2), unless the *z*-CAIPI shift is a factor of *R_z_*: in that case the shot-selective *y*-blip is only *R_z_*/|*Δz*| times larger (Fig. 1 C, row 2).

## Methods

Segmentation can be applied along the shot-selective CAIPI trajectory to achieve even larger *y*-blips as a multiple of the shot-selective *y*-blip (6). In this work, we go one step further and propose segmentation along the blipped-CAIPI trajectory to also obtain all intermediate *y*-blips and *EFs*. This segmented k-space blipped-CAIPI (skipped-CAIPI) approach maximally unlinks the *y*-blip from the CAIPI pattern. Skipped-CAIPI is demonstrated on the example of a 3D-EPI sequence. However, it can be applied to SMS-3D space w.l.o.g. (38–40).

### Skipped-CAIPI

A skipped-CAIPI sampling trajectory is defined by the CAIPI pattern and a segmentation factor, *S* ≥ 1. Along the blipped-CAIPI trajectory, *S* − 1 echoes are skipped and acquired in subsequent shots (Fig. 1, row 3). Analogous to segmented EPI, the *y*-blip becomes *S · R_y_*. In fact, traditional segmented EPI corresponds to skipped-CAIPI with *Δz* = 0 (Fig. 1 A). Blipped-CAIPI corresponds to skipped-CAIPI with *Δz* ≠ 0 and *S* = 1 (Fig. 1 B-C, row 1). The CAIPI pattern is not affected by the segmentation factor. For the sampling trajectory, we use the shorthand notation:

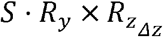

Like blipped-CAIPI, most skipped-CAIPI trajectories imply *z*-blips between echoes. Two *z*-blips with opposed polarity are repeated in a sequential order with a certain cycle. The *z*-blip cycle of blipped-CAIPI is *n*_1_ = *R_z_* (e.g. Fig. 1 B, row 1), unless *Δz* is a factor of *R_z_*, which results in *n*_1_ = *R_z_*/|*Δz*| (e.g. Fig. 1 C, row 1). Consequently, employing skipped-CAIPI with *S* = *n*_1_ equals shot-selective CAIPI (Fig. 1 B-C, row 2). Although *S* = 2*n*_1_, 3*n*_1_,… trajectories may be considered “segmented shot-selective”, we refer to them as “skipped-CAIPI without *z*-blips” to avoid ambiguity. For more details, we refer the reader to the Appendix.

Like segmented EPI, skipped-CAIPI hypothetically works for all *S* up to *N_y_*/*R_y_*, even though very large *S* are not efficient. Given a CAIPI pattern, 1/*n*_1_ of all corresponding trajectories do not imply *z*-blips, including shot-selective CAIPI (*S* = *n*_1_). An equal or greater number, (*n*_1_ − 1)/*n*_1_ of all trajectories, implies *z*-blips, including blipped-CAIPI (*S* = 1). See Supporting Information Figures S1 for a plot of *EF* vs. *y*-blip, and S2 for a graphical representation of all sampling trajectories with identical *z*-blips for the same three CAIPI pattern examples. Vice versa, if the *y*-blip shall have a fixed value, Supporting Information Figures S3-5 show all possible CAIPI patterns up to *R* = 8 according to (37) and sampling trajectories that yield a *y*-blip of 2, 3 and 4.

### Experiments

All experiments were performed on a 3T MAGNETOM Skyra (Siemens Healthineers, Software version VE11C) equipped with a 45mT/m, 200T/m/s gradient system and a 32 channel head coil. A custom RF-spoiled 3D-EPI sequence with echo time shifting was used (Fig. 1 D-E).

One healthy male subject was scanned after providing informed consent in accordance with local institutional review board regulations. Two protocol types tailored towards whole-brain functional MRI (Tab. 1), and rapid T_1_-weighted structural imaging were used (Tab. 2). All images were reconstructed using a generic vendor-provided GRAPPA (41) implementation compatible with 2D CAIPIRINHA (“IcePAT”).

**Table 1:** Sampling trajectories and sequence parameters for time series EPI acquisitions.

| R | y-blip: | $S \cdot R_y = 2$ ($N_{shots} / TR_{vol}$) | $S \cdot R_y = 3$ ($N_{shots} / TR_{vol}$) | $S \cdot R_y = 4$ ($N_{shots} / TR_{vol}$) |
| --- | --- | --- | --- | --- |
| 4 | Blipped | $1 \cdot 2 \times 2_{z1}$ (36 / 1.90s) | N/A | N/A |
| | Shot-s. | $2 \cdot 1 \times 4_{z2}$ (36 / 1.90s) | N/A | $2 \cdot 2 \times 2_{z1}$ (80 / 4.70s) |
| | Skipped | $2 \cdot 1 \times 4_{z1}$ (36 / 1.90s) | $3 \cdot 1 \times 4_{z2}$ (60 / 3.5s) | $4 \cdot 1 \times 4_{z2}$ (80 / 4.70s) |
| 5 | Skipped | $2 \cdot 1 \times 5_{z2}$ (28 / 1.48s) | $3 \cdot 1 \times 5_{z2}$ (48 / 2.8s) | $4 \cdot 1 \times 5_{z2}$ (64 / 3.80s) |
| 6 | Blipped | $1 \cdot 2 \times 3_{z1}$ (24 / 1.26s) | $1 \cdot 3 \times 2_{z1}$ (42 / 2.5s) | N/A |
| | Shot-s. | $2 \cdot 1 \times 6_{z3}$ (24 / 1.27s) | $3 \cdot 1 \times 6_{z2}$ (42 / 2.5s) | N/A |
| | Skipped | $2 \cdot 1 \times 6_{z2}$ (24 / 1.28s) | $3 \cdot 1 \times 6_{z3}$ (42 / 2.5s) | $4 \cdot 1 \times 6_{z2}$ (56 / 3.30s) |
| 7 | Skipped | $2 \cdot 1 \times 7_{z2}$ (20 / 1.07s) | $3 \cdot 1 \times 7_{z2}$ (36 / 2.1s) | $4 \cdot 1 \times 7_{z2}$ (48 / 2.90s) |
| 8 | Blipped | $1 \cdot 2 \times 4_{z2}$ (18 / 0.95s) | N/A | $1 \cdot 4 \times 2_{z1}$ (40 / 2.40s) |
| | Shot-s. | $2 \cdot 1 \times 8_{z4}$ (18 / 0.95s) | N/A | $2 \cdot 2 \times 4_{z2}$ (40 / 2.40s) |
| | Skipped | $2 \cdot 1 \times 8_{z2}$ (18 / 0.97s) | $3 \cdot 1 \times 8_{z2}$ (30 / 1.75s) | $4 \cdot 1 \times 8_{z3}$ (40 / 2.40s) |
|  | EF: | 54 | 48 | 45 |
|  | Res.: | 2 mm isotropic | 1.5 mm isotropic | 1.2 mm isotropic |
| | Matrix: | $108 \times 108$ | $144 \times 144$ | $180 \times 180$ |
| | $N_{slices}$: | 72 (70 for R=5,7) | 80 (84 for R=6,7) | 80 (84 for R=6,7) |
| | $BW_x$: | 1714 Hz/pixel | 1240 Hz/pixel | 1112 Hz/pixel |
|  | ESP<sup>a</sup>: | 0.71-0.75 ms | 0.95-0.99 ms | 1.02-1.10 ms |
| | $BW_y$<sup>a</sup>: | 24.7-26.1 Hz/pixel | 21.0-21.9 Hz/pixel | 20.2-21.8 Hz/pixel |
|  | TR<sup>a</sup>: | 52.8-53.9 ms | 58.3-59.5 ms | 58.8-60.4 ms |
| | $N_{vol}$: | 55 (1-5 removed for tSNR) | 45 (1-3 removed for tSNR) | 35 (1-2 removed for tSNR) |
|  | Fig. 2 | A | B | C |

<sup>a</sup> ESP varies at constant $BW_x$ due to varying z-blip durations (varying ramp sampling). $N_{shots} = S \cdot N_{slices} / R_z$ : number of shots per volume measurement. $TR_{vol}$ : acquisition time per volume measurement. EF: EPI factor. Res.: resolution. $N_{slices}$ : number of slices. $BW_x$ : readout bandwidth. ESP: echo spacing. $BW_y$ : phase encode bandwidth. TR: repetition time per shot. $N_{vol}$ : number of volume measurements.

### Functional MRI protocol type

Time series data were acquired with 26 different protocols according to Tab. 1 using oblique axial slice orientation. In terms of temporal SNR (tSNR), the selected protocols corresponded to the best applicable blipped-, shot-selective and skipped-CAIPI trajectories with 4 ≤ *R* ≤ 8. As *ESP* increases with resolution, decreasing *EFs* of 54, 48 and 45 were necessary to achieve TE=30ms, which corresponds to *y*-blips 2, 3 and 4. A 15° binomial-121 water excitation pulse was employed to maximize gray matter signal (sinc, bandwidth - time-product=30, slab scaled to 90% of nominal slice FOV). Between each excitation and EPI readout a standard integrated phase correction (PC) scan was acquired.

### Dual-polarity ACS

As part of each time series acquisition, 48×48 initial autocalibration scan (ACS) lines prepared by 200 steady-state dummy shots were acquired using a (*S · R_y_*) · 1 × 1_*z*0_ segmented EPI sampling (*BW_y_* matched to subsequent imaging) with minimal TE and TR (Fig. 1 D). Every shot was acquired twice with alternating readout polarity, each echo phase corrected (almost ghost-free) and then complex averaged (completely ghost-free). This simplified dual-polarity ACS corresponds to the first steps of dual-polarity GRAPPA processing (42) and integrates easily without reconstruction customization.

### Structural MRI protocol type

All structural scans according to Tab. 2 used sagittal slice orientation (readout: head-feet). All EPI scans employed a single hard pulse excitation set to approximately 2.4ms duration for water excitation at 3T (6,43). A single external PC scan (5° excitation) followed by 100 FLASH dummy shots and ACS lines (44) were acquired as a prescan (Fig. 1 E). The total duration, *T_pre_*, was included in all stated TAs.

**Table 2:** Sampling trajectories and sequence parameters for structural EPI acquisitions.

| y-blip: | S·R<sub>y</sub>=21, 22, 21 (TR<sub>vol</sub> / TA) | S·R<sub>y</sub>=30 (TR<sub>vol</sub> / TA) | S·R<sub>y</sub>=30, 20, 10 (TR<sub>vol</sub> / TA) |
| --- | --- | --- | --- |
| Skipped | 21 · 1 × 2<sub>z1</sub> (26s / 0:31)<br>11 · 2 × 2<sub>z1</sub> (14s / 0:19)<br>7 · 3 × 2<sub>z1</sub> (10s / 0:14) | 30 · 1 × 5<sub>z2</sub> (30.0s / 0:44)<br>10 · 3 × 2<sub>z1</sub> (25.0s / 0:39)<br>30 · 1 × 7<sub>z3</sub> (21.0s / 0:35) | 15 · 2 × 2<sub>z1</sub> (37.5s / 0:52-2:44<sup>a</sup>)<br>10 · 2 × 2<sub>z1</sub> (31.0s / 0:45)<br>5 · 2 × 2<sub>z1</sub> (24.5s / 0:39) |
| EF: | 7 | 9 | 9, 14, 27 |
| Res.: | 1.2 mm isotropic | 0.8 mm isotropic | 0.8 mm isotropic |
| Matrix: | 192 × 192 | 270 × 270 | 270 × 270 |
| FOV: | 230 mm × 230 mm | 216 mm × 216 mm | 216 mm × 216 mm |
| N<sub>slices</sub>: | 160 | 200 (196 for R=7) | 200 |
| P. Fourier | 6/8 × 1 | 1 × 1 | 1 × 1 |
| TE: | 4.35ms | 10 ms | 10.0, 12.8, 22.2 ms |
| TR: | 15.7ms | 25 ms | 25, 31, 48 ms |
| BW<sub>x</sub>: | 1184 Hz/pixel | 806 Hz/pixel | 806 Hz/pixel |
| ESP: | 1.07, 1.09, 1.07 ms | 1.55 ms | 1.55, 1.51, 1.46 ms |
| BW<sub>y</sub>: | 102.2, 105.1, 102.2 Hz/pixel | 71.7 Hz/pixel | 71.7, 49.1, 25.3 Hz/pixel |
| ACS | 48 × 16 | 48 × 48 | 48 × 48 |
| T<sub>pre</sub> in TA: | 4.9 s | 13.9 s | 13.9 s |
| Fig. 3 | A | B | C, D<sup>a</sup> |

<sup>a</sup> N<sub>vol</sub>=4 volume measurements. TR<sub>vol</sub>: acquisition time per volume measurement. TA: total acquisition time including multiple volume measurements and preparation time. EF: EPI factor. Res.: resolution. N<sub>slices</sub>: number of slices. P. Fourier: partial Fourier factor. TE: echo time. TR: repetition time per shot. BW<sub>x</sub>: readout bandwidth. ESP: echo spacing. BW<sub>y</sub>: phase encode bandwidth. ACS: autocalibration scan (y × z matrix). T<sub>pre</sub>: preparation time (external PC + ACS dummy shots + FLASH ACS).

For submillimeter scans, TR=25ms and a flip angle (FA) of 30° was used. Four volumes were acquired using a 15 · 2 × 2_*z*1_ sampling for retrospective averaging with anterior-posterior primary phase encode direction (AP). The scan was repeated once with inverted phase encode direction (PA). As an EPI factor-free reference, the vendor-provided multiecho (ME) FLASH sequence, lacking CAIPIRINHA sampling, was used with otherwise identical parameters, except for a slightly altered FOV (230mm along RO, 192 slices), 0.1ms hard pulse excitation and readout BW_x_=380Hz/pixel for each of six bipolar echoes (TE∈[3.8,l9.3)ms). The resulting TA=5:36 approximates the combined duration of AP and PA EPI scans with four averages each (5:28).

Additional TR/TE/FA/*EF*-matched EPIs with alternative CAIPI patterns were acquired. Furthermore, the 2 × 2_*z*1_ pattern was sampled with varying segmentation factors, *S* = 15,10,5. Finally, 1.2mm isotropic data were acquired, adapting and extending a rapid T_1_-weighted scan recently proposed for clinical use (6): for consistent TR=15.7ms and TE=4.35ms across different CAIPI patterns, *EF* = 7 and minimal sequence dead times and shots per measurement were maintained by adjusting *S* as facilitated by skipped-CAIPI (Tab. 2).

### Processing

All processing was performed using FSL software (45) and python. Time series were preprocessed using FEAT, including temporal high-pass filtering (100s) and motion correction. 5, 3 or 2 initial volumes were discarded before computing voxel-wise tSNR, finally divided by 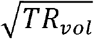 to assess tSNR efficiency.

The combined eight volumes of the anatomical AP and PA 15 · 2 × 2_*z*1_ scans were processed using TOPUP to estimate and correct for motion and geometric distortions. The corrected magnitude images were averaged for visual comparison to the ME-FLASH image. For the latter, all TE magnitude images were averaged for maximal SNR and comparable T_2_*-weighting.

## Results

Fig. 2 shows a sagittal example view of the best time series tSNR and efficiency results for *R* = 4,5,6,7,8 at isotropic resolutions of 2mm (A), 1.5mm (B) and 1.2mm (C). The respective volume TRs and the quartiles of the whole-brain tSNR efficiencies are printed above and below the maps, respectively. Results of the alternative protocols according to Tab. 1, for which the CAIPI pattern resulted in minor-to-major efficiency degradation for the same *R*, are shown in Supporting Information Figure S6.

**Figure 2:**
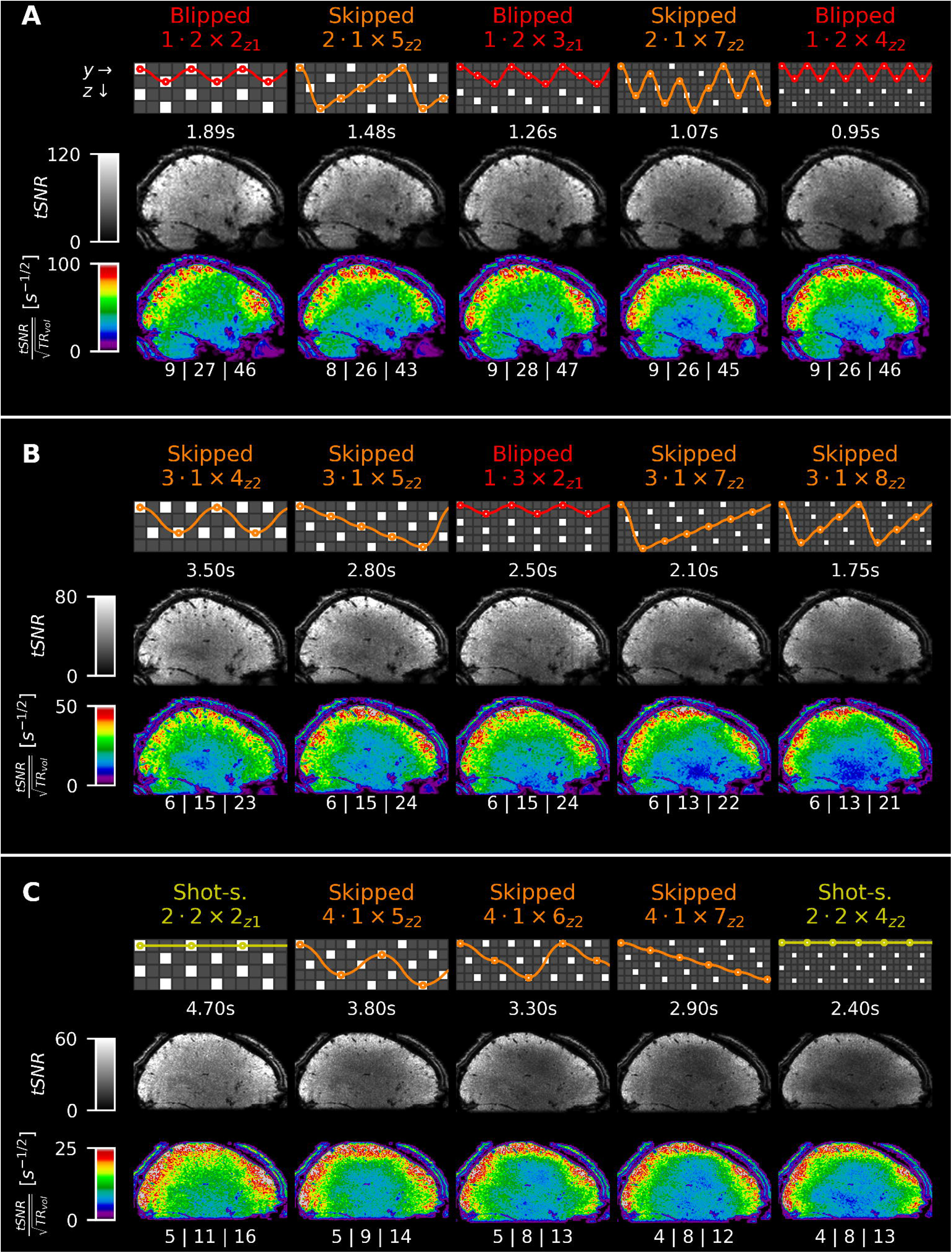
Temporal SNR maps and tSNR efficiency maps for the time series data acquired with different CAIPI patterns resulting in R = 4 – 8 (from left to right), which provided the best overall tSNR efficiency at 2.0 mm (A, y-blip 2), 1.5 mm (B, y-blip 3) and 1.2 mm isotropic resolution (C, y-blip 4) under the described experimental conditions. Out of 15 presented cases, 9 were not applicable previously (labeled “Skipped”). The respective CAIPI pattern along with one example trajectory and TR_vol_ is indicated above each tSNR map. Whole-brain tSNR efficiency quartiles are indicated below the maps. For the remaining, second-to-third best results according to alternative CAIPI scans of Tab. 1, as well as example magnitude images, see Supporting Information Figure S6.

The first three columns of Fig. 3 show axial example views of single volume T_1_-weighted EPI scans at 1.2mm isotropic resolution according to (6) with varying CAIPI patterns and constant *EF* = 7 (A), and at 0.8mm isotropic resolution with varying CAIPI patterns and constant *EF* = 9 (B), and with varying *EF* and constant 2 × 2_*z*1_ CAIPI pattern (C). The two rightmost columns show corresponding views and magnified zooms following magnitude averaging (D, from top to bottom): 4 volumes, 8 volumes following motion and distortion correction, and 6 TEs of the FA/TR/TA-matched ME-FLASH. Yellow arrows indicate the effect of distortion correction. Supporting Information Figure S7 shows corresponding orthogonal views and additional examples with local SNR estimates and an MP-RAGE scan for anatomical reference. Supporting Information Figure S8 shows the actual k-space samplings used.

**Figure 3:**
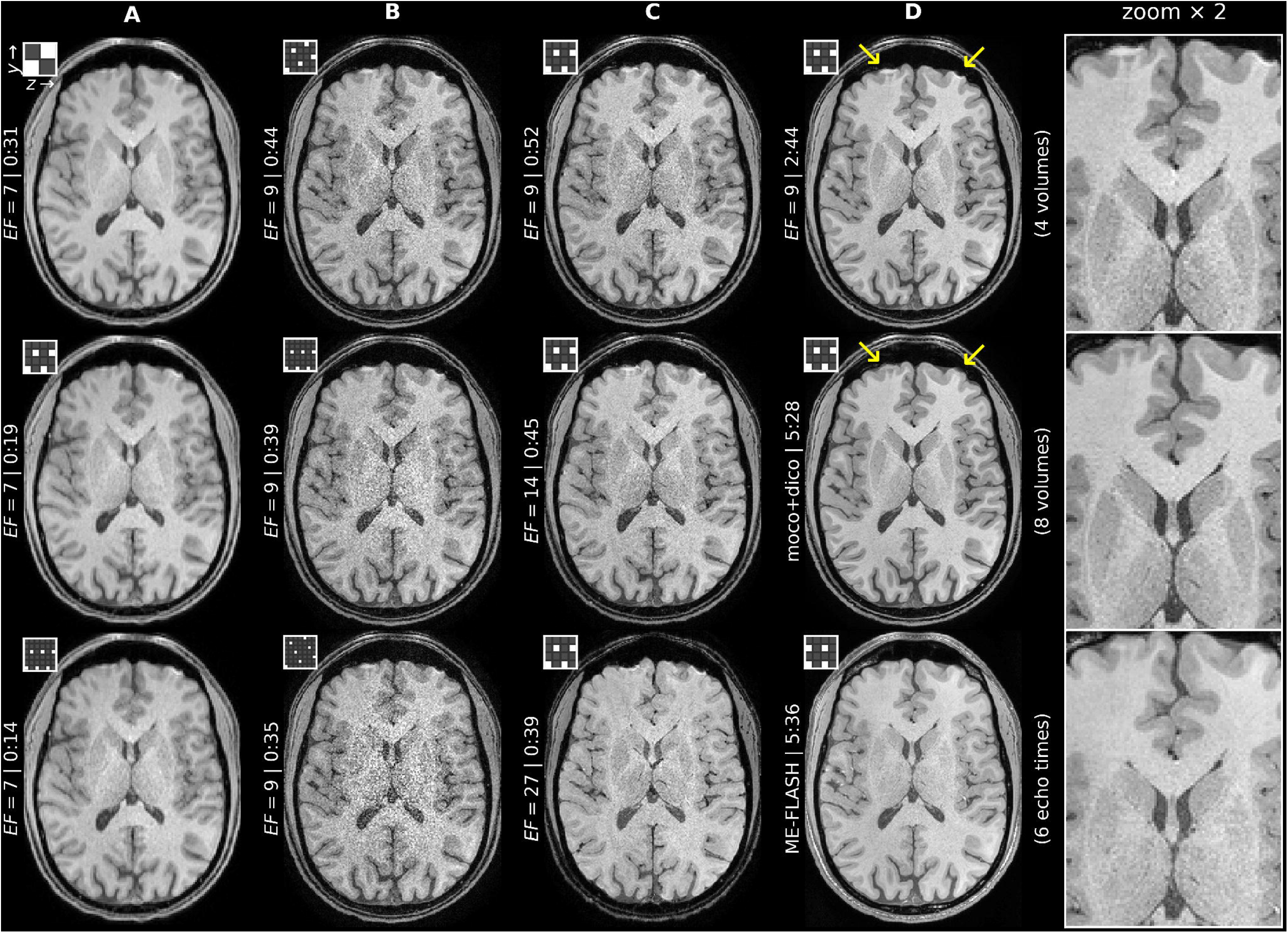
Axial example slice of several rapid T_1_-weighted high-resolution whole-head scans (TA stated in [min:s]) along with the respective elementary CAIPIRINHA sampling cells (37). A: 1.2 mm isotropic protocols adapted from (6) (row 1), with increasing CAIPI undersampling whilst keeping EF = 7 constant B-D: 0.8 mm isotropic protocols with increasing CAIPI undersampling whilst keeping EF = 7 constant (B), with increasing EF whilst keeping CAIPI pattern constant (C), and four- and eight-fold magnitude-average without and with correction for motion and geometric distortions compared to the echo-average of a TR- and TA-matched ME-FLASH (D). See Supporting Information Figure S7 for orthogonal example slices of (D), additional scan comparisons and local SNR estimates. See Supporting Information Figure S8, for complete k-space sampling of scans (A-C). moco: motion correction, dico: distortion correction. ME-FLASH: multi-echo FLASH. EF: EPI factor.

## Discussion

Previous blipped-CAIPI and shot-selective CAIPI approaches corresponded to exactly two possible EPI trajectories given a specific CAIPI pattern. By implication, instead of selecting the optimal CAIPI pattern, it often had to be selected according to the associated *y*-blip that fit the matrix size, TE, TR, or *EF* prerequisites, in particular at high temporal resolution applications such as fMRI. Large *y*-blips for high spatial resolution with minimal geometric distortions require segmentation, which previously only provided limited CAIPI options without *z*-blips. In both resolution domains, skipped-CAIPI increases protocol flexibility without compromising controlled aliasing quality. In fact, by selecting a g-factor optimal CAIPI pattern first, and then selecting a segmentation factor to adjust *EF* or *BW_y_* via the *y*-blip, chances are relatively high (1:*n*_1_) to yield a trajectory without *z*-blips, but *n*_1_ − 1 times higher to yield a trajectory with *z*-blips (Supporting Information Figures S1-2). From this point of view, skipped-CAIPI may as well be understood as segmented EPI with a CAIPI shift. The segmentation factor blends between the special cases of blipped-CAIPI (*S* = 1), shot-selective CAIPI (*S* = *n*_1_) and skipped-CAIPI without *z*-blips (*S* = 2*n*_1_, 3*n*_1_,…).

The resulting flexibility is demonstrated by the T_1_-weighted 3D-EPI examples of Fig. 3, where one parameter was varied while others were kept fix without introducing unnecessary sequence dead times, as facilitated by skipped-CAIPI. FLASH-like acquisitions with sufficient contrast-to-noise ratio (e.g. 1.2mm data, A) can be used for rapid MR parameter quantification (10,46–48) and they also have great clinical potential as a rapid structural imaging alternative (6), especially if extended to other contrasts (4,49). At 7T, an established method for efficient high-resolution T_2_*/susceptibility-weighted imaging uses segmented 3D-EPI without CAIPI sampling (1). In this example, skipped-CAIPI could either reduce the g-factor at unchanged undersampling factor of 2, or accelerate further with negligible g-factor increase. Either way, TE, TR and *EF* could be left unchanged. Reduced intra-volume motion sensitivity, as single measurements get faster, and the option of intervolume correction for motion and distortion before averaging pose a general advantage over a single 3D-(ME)-FLASH scan of the same total acquisition and spatial coverage. This is demonstrated by Fig. 3 (D) and Supporting Information Figure S7. Although the subject was instructed to lie still, the 8-fold EPI average even shows sharper image details at similar SNR. More SNR may be gained, if corrections (including voxel-wise phase drift) and averaging are performed on the complex images. In terms of SNR, complex image averaging is equivalent to k-space oversampling (or extended slab coverage (1)) as long as the g-factor is negligible. Therefore: the ability to acquire faster without reducing SNR efficiency (e.g. EPI vs. FLASH) or to increase SNR without slowing down acquisition (e.g. segmented EPI with CAIPI vs. without CAIPI) reflects one strength of skipped-CAIPI.

Aside from high-resolution imaging, skipped-CAIPI is also beneficial, if smaller *y*-blips are required, e.g. for fMRI. According to Supporting Information Figures S3-5, even with small *y*-blips, the majority of CAIPI patterns could previously not be sampled by means of EPI. For several blipped-, shot-selective and skipped-CAIPI trajectories, the combined effect of *g, R*, receive sensitivity, physiological noise (50) and temporal resolution was assessed by tSNR and tSNR efficiency. The free choice of CAIPI pattern, independent of the required *y*-blip, has demonstrated maximum efficiency across previously inaccessible undersampling factors and resolutions (Fig. 2). For instance, *y*-blip 3 scans can now additionally be acquired using well-suitable *R* = 2,4*, 5*, 7*, 8*, 10, etc. CAIPI patterns compared to previously only *R* = 3,6*, 9, etc. (*demonstrated here). Note: which CAIPI pattern is optimal may differ when using a different array coil, slice orientation or slice coverage. A limitation of any EPI sampling is that the *y*-blip is a multiple of *R_y_*. For instance, the 2 × 2_z1_ CAIPI pattern used for the anatomical protocol (*y*-blip 30) could not be used for the 1.5mm functional protocol (*y*-blip 3). On the other hand, *R_y_* = 1 skipped-CAIPI trajectories, such as *S* · 1 × 4_z2_, are generally applicable and can be beneficial in terms of g-factor. Therefore: filling gaps of previously inaccessible undersampling options with minimal g-factor penalty reflects another strength of skipped-CAIPI.

Although skipped-CAIPI has been discussed for the 2D phase encoding space of 3D-EPI, it can just as well be applied to the SMS-3D sampling space (38,39). Shot-selective CAIPI, for instance, has already been used with SMS-EPI for dynamic contrast-enhanced imaging (5). This implementation could be adopted for skipped-CAIPI sampling, either without *z*-blips or with *z*-blips. A skipped-CAIPI SMS option may be advantageous for certain applications, or physiological noise versus thermal noise regimes (51). However, the limitations of traditional segmented EPI compared to single-shot EPI, i.e. an increased sensitivity to shot-to-shot signal variations, also apply to SMS-EPI using skipped-CAIPI or shot-selective CAIPI sampling. On the other hand, high-resolution MRI, which may benefit most from skipped-CAIPI sampling, also benefits most from volumetric (3D) imaging (33,52). For multi-echo fMRI (53–56) or ultra-high-field fMRI, where often partial Fourier sampling and large *R_y_* are employed to minimize *EF*, TE and geometric distortions, alternative skipped-CAIPI sampling is valuable, in particular with high-density receive arrays (13,18) or for laminar resolution imaging (57,58). Furthermore, eased matching of the phase encode bandwidth between scans with different matrix sizes or CAIPI patterns fosters rapid, distortion-matched high-contrast acquisitions (19,59,60) even at resolutions differing from the corresponding functional scan.

## Conclusion

Introducing segmentation into blipped-CAIPI - or equivalently, introducing a CAIPI shift into segmented EPI - allows to select a g-factor-optimal 2D-CAIPIRINHA pattern and the primary phase encode gradient blip independently to adjust geometric distortions, echo times, repetition times, etc. Almost arbitrary EPI factors combined with minimal parallel imaging noise are thus made available to numerous MR imaging methods, including volumetric high-resolution imaging. While the implementation is simple, the versatility gain of typical and non-typical applications of 3D-EPI or SMS-EPI can be vast.

## Appendix

### Z-blips

The magnitudes of the two *z*-blips, 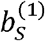 and 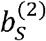, of any skipped-CAIPI trajectory are determined by *S* ≥ 1 and the CAIPI parameters (in *z*-notation) *R_z_* > 1 and 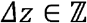:

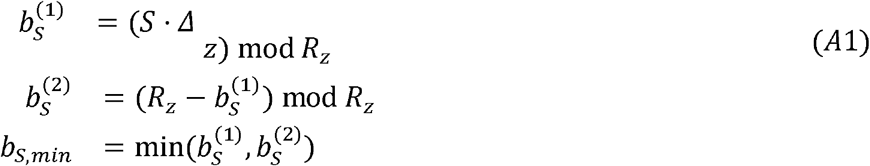

Here, mod denotes the remainder of the division operator (with the sign of the divisor).

### Z-blip cycle

Every *n_s_* echoes, the sequential *z*-blip order repeats:

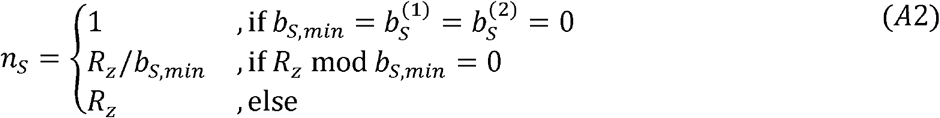

The same *z*-blips/cycle apply for all segmentation factors *S*′ = *S* + *m* · *n*_1_ ≥ 1 with 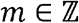:

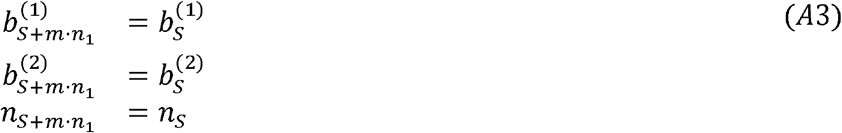

Here, *n*_1_ is the blipped-CAIPI *z*-blip cycle. The special cases *S* = *n*_1_ and *S*′ = 2*n*_1_, 3*n*_1_,… imply no *z*-blips (cf. Supporting Information Figure S2).

Note: if *Δz* ∈ [−└*R_z_*/2┘, └*R_z_*/2┘]\{0}, as assumed in this work, for blipped-CAIPI Eq. A2 simplifies to

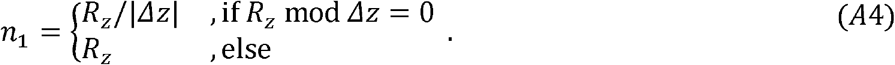

## Supporting Information

***Supporting Information Figure S1**: EPI factors (EF) as a result of segmentation when combining a relatively small (N_y_ = 108) and large (N_y_ = 270) phase encode matrix size with three different CAIPI pattern examples. The phase encode bandwidth is proportional to the y-blip (horizontal axis). A/B: logarithmic/linear EF axis (according to a reduced range of segmentation factors indicated by dotted rectangle in A). Using traditional segmented EPI (blue line: same undersampling pattern, but without CAIPI shift), a wide range of EPI factors and y-blips is applicable. Blipped-CAIPI and shot-selective CAIPI only correspond to two distinct EPI factors and y-blips. With skipped-CAIPI, all EPI factors and y-blips of segmented EPI can be applied with a reduced g-factor. Inset displays in B show the respective blipped-CAIPI and shot-selective CAIPI trajectories and the first unique skipped-CAIPI option (first shot: thick, bright curve; subsequent shots: thinner, darker curves). Vertical grid lines in B indicate EPI trajectories without z-blips (S = n_1_, 2n_1_, 3n_1_,…, where n_1_: blipped-CAIPI z-blip cycle)*.

***Supporting Information Figure S2:** Z-blips and z-blip cycles of various CAIPI trajectories for the example patterns of S1. For the sake of generality, 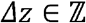 (see Appendix). Both trajectories for negative and positive CAIPI shifts are indicated on the example of the first of S shots. Identical z-blips/cycle apply to all following shots (albeit starting at a different offset in the cycle). Identical z-blips/cycle also occur for the same CAIPI pattern with segmentation factors S′ = S,S + n_1_, S + 2n_1_,…, whereby n_1_ denotes the blipped-CAIPI z-blip cycle (S = 1) associated to the CAIPI pattern*.

***Supporting Information Figure S3:** Applicable and not applicable EPI R ≤ 8 undersampling with a fixed y-blip S · R_y_ = 2. Only Δ_z_ ≥ 0 patterns are plotted, but corresponding −Δ_z_ patterns are counted, if applicable (±). First trajectory/subsequent trajectories indicated by thick and bright/thinner and darker curves. Sampling trajectories used for experimental time series acquisition in this work indicated by * (cf. Tab. 1, Fig. 2, Supporting Information Figure S6)*.

***Supporting Information Figure S4:** Same as S3 with fixed y-blip S · R_y_ = 3*.

***Supporting Information Figure S5:** Same as S3 with fixed y-blip S · R_y_ = 4. The R = 2,4,6,8 skipped-CAIPI schemes include trajectories without z-blips*.

***Supporting Information Figure S6:** Temporal SNR and tSNR efficiency maps of second-to-third best time series data acquired in this work. Even though these were the best applicable blipped-CAIPI, shot-selective CAIPI and skipped-CAIPI trajectories for the respective total undersampling factors at fixed y-blips of 2 (A, 2 mm isotropic), 3 (B, 1.5mm isotropic) and 4 (C, 1.2mm isotropic), the corresponding CAIPI patterns lead to marginally-to-substantially worse results than the optimal CAIPI patterns shown in Fig. 2. For instance, skipped-CAIPI 2 · 1 × 6_z2_ performed only marginally worse than the optimal blipped-CAIPI 1 · 2 × 3_z1_ trajectory (cf. Fig. 2). The 1 × 6_z3_ CAIPI pattern performed particularly poorly, irrespective of the trajectory used (cf. A,B). However, the performance may differ using a different coil, slice coverage or slice orientation. The enlarged views on the right show the mean magnitude images of the respective R = 8 skipped-CAIPI acquisition. The 1.5mm example is also shown without automatic receive sensitivity correction (“prescan normalize”) to indicate the receive sensitivity*.

***Supporting Information Figure S7:** Orthogonal example slices of T_1_-weighted high-resolution whole-head scans (TA stated in [min:s]) corresponding to the axial slices shown in Fig. 3 D and three additional examples (right panel). Eight magnitude-averages of the 15 · 2 × 2_z1_ EPI scan (b) show improved anatomical accuracy due to motion and distortion correction (moco+dico, yellow arrows) compared to four averages without corrections (a), and even improved resolution of details compared to the echo-average of a TR/FA/TA-matched ME-FLASH (c, red arrows). *A single-echo (TE=3.8ms) FLASH scan obtain with reduced TR=12ms and FA=20° but otherwise identical parameters (d) and two magnitude-averages of the 15 · 2 × 2_z1_ EPI following moco+dico (e) are additionally shown. Detailed views with a zoom factor of two are shown below all example slices. Blue numbers indicate rough SNR estimates from squared white matter regions indicated in the zoomed axial view of the corresponding MP-RAGE (f 0.9mm isotropic, 240mm×240mm FOV, 192 slices, TE=2.32ms, TR=2.3s, TI=900ms, flip angle 8°, 2 × 1 GRAPPA, 24 integrated ACS lines, TA=5:21)*.

***Supporting Information Figure S8:** Visualization of the skipped-CAIPI samplings used for high-resolution T_1_-weighted 3D-EPI scans in this work (cf. Tab. 2, Fig. 3, Supporting Information Figure S7) along with the trajectory that covers the k-space center echo (defining the image echo time, TE). Complete and zoomed k-space views use the echo number (“Echoes”) and the actual echo time (“Echo times (zoom)”) for color-coding, respectively. Without echo time shifting (delaying the start time of each EPI trajectory by a fraction of the echo spacing according to the first primary phase encode value of the EPI trajectory, cf. Fig. 1), the echo times color-coding would resemble the step-like echoes color coding. Image artifacts would arise, where local magnetic field inhomogeneities are strong. However, due to echo time shifting, the echo times and the MR signal magnitude and phase at each frequency-encoded echo center evolve continuously along the primary phase encode direction (31,32). As skipped-CAIPI can be understood as traditional segmented EPI with a CAIPI shift, this applies with or without CAIPI shift*.

