## Supporting Information Figures S1-8 for "Segmented K-Space Blipped-Controlled Aliasing in Parallel Imaging (Skipped-CAIPI) for High Spatiotemporal Resolution Echo Planar Imaging"

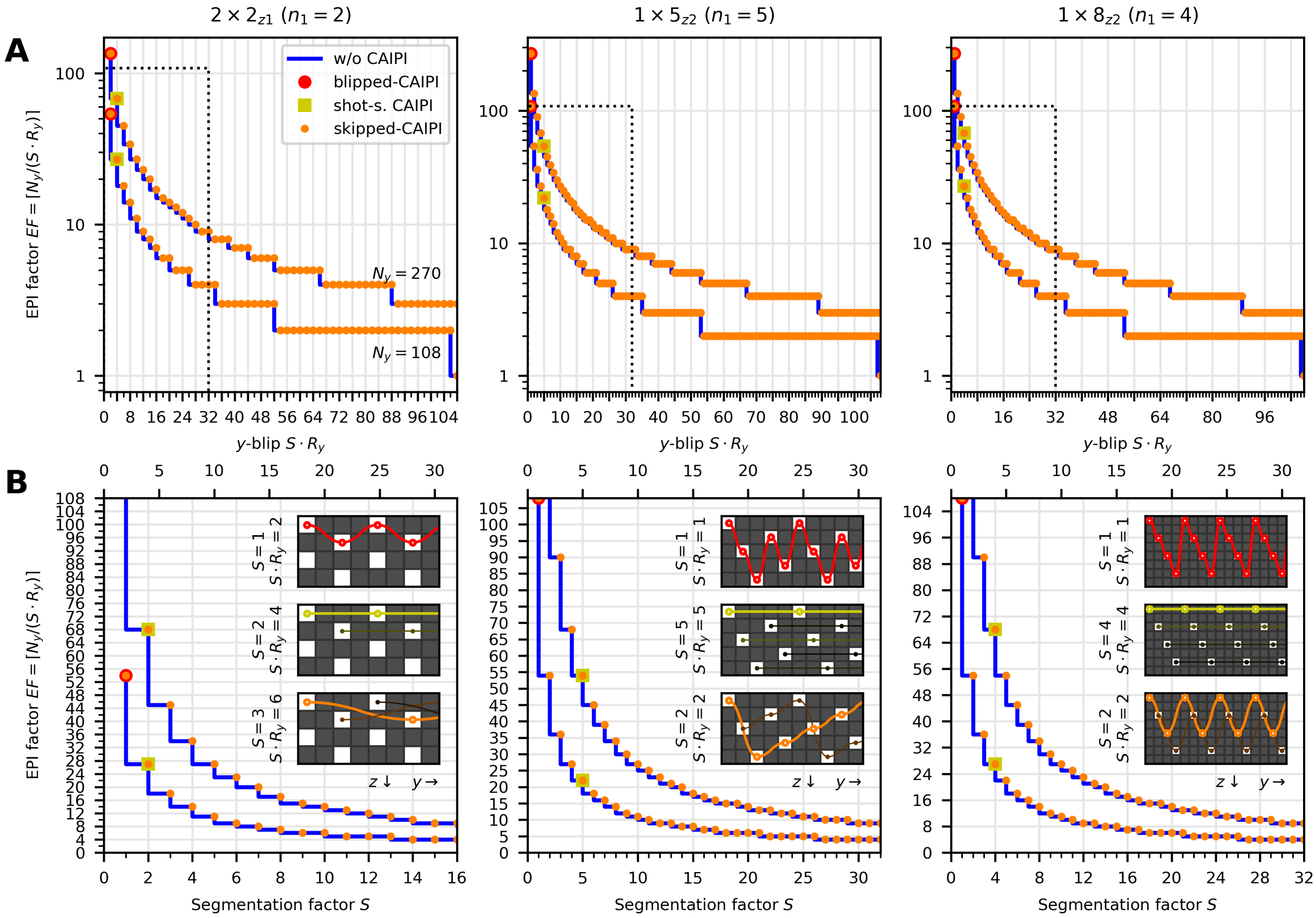


**Supporting Information Figure S1:** EPI factors ($EF$) as a result of segmentation when combining a relatively small ($N_y$ = 108) and large ($N_y$ = 270) phase encode matrix size with three different CAIPI pattern examples. The phase encode bandwidth is proportional to the $y$-blip (horizontal axis). A/B: logarithmic/linear $EF$ axis (according to a reduced range of segmentation factors indicated by dotted rectangle in A). Using traditional segmented EPI (blue line: same undersampling pattern, but without CAIPI shift), a wide range of EPI factors and $y$-blips is applicable. Blipped-CAIPI and shot-selective CAIPI only correspond to two distinct EPI factors and $y$-blips. With skipped-CAIPI, all EPI factors and $y$-blips of segmented EPI can be applied with a reduced g-factor. Inset displays in B show the respective blipped-CAIPI and shot-selective CAIPI trajectories and the first unique skipped-CAIPI option (first shot: thick, bright curve; subsequent shots: thinner, darker curves). Vertical grid lines in B indicate EPI trajectories without $z$-blips ($S = n_1, 2n_1, 3n_1, ...$, where $n_1$: blipped-CAIPI $z$-blip cycle).

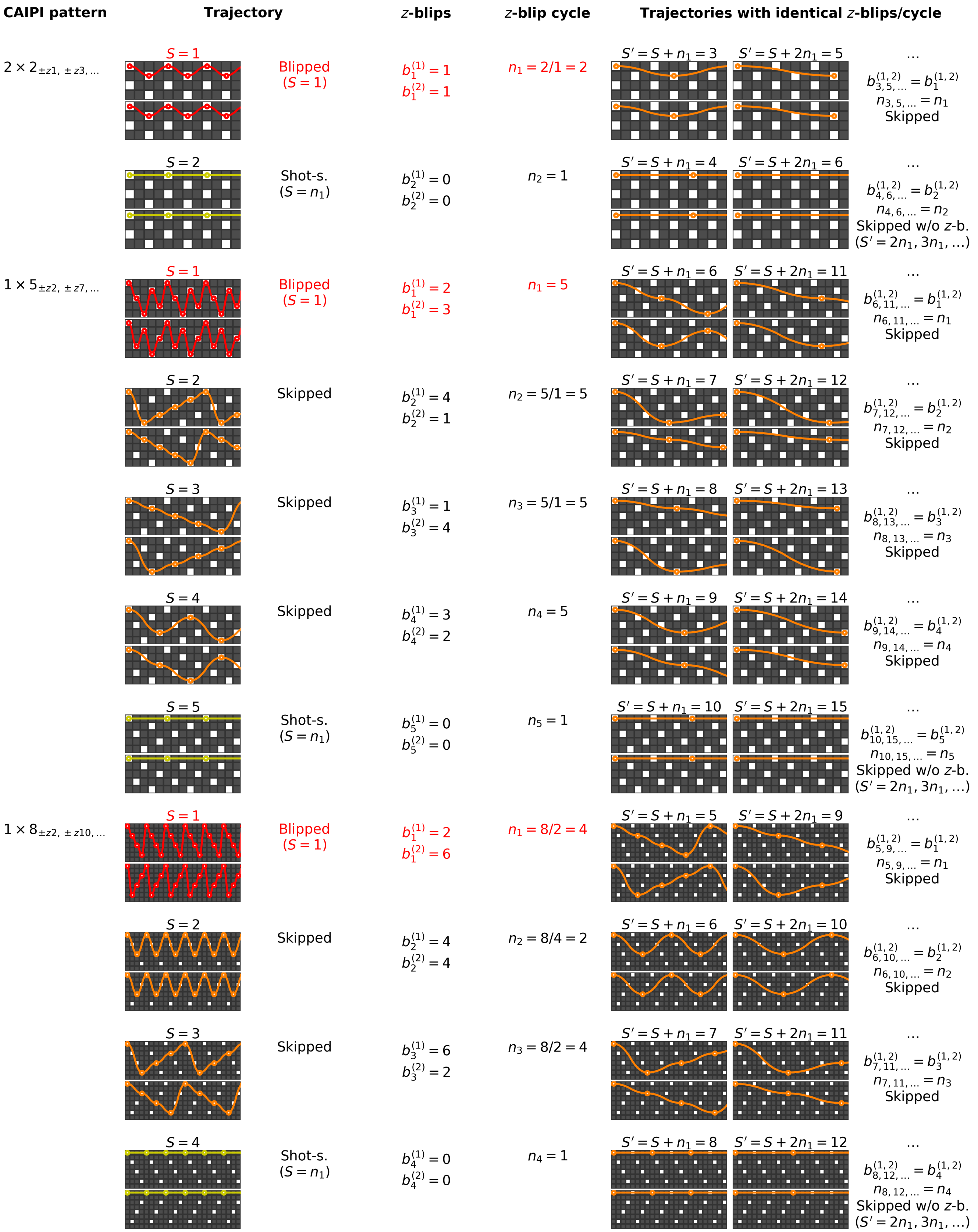


**Supporting Information Figure S2:** $Z$-blips and $z$-blip cycles of various CAIPI trajectories for the example patterns of S1. For the sake of generality, $\Delta z \in \mathbb{Z}$ (see Appendix). Both trajectories for negative and positive CAIPI shifts are indicated on the example of the first of $S$ shots. Identical $z$-blips/cycle apply to all following shots (albeit starting at a different offset in the cycle). Identical $z$-blips/cycle also occur for the same CAIPI pattern with segmentation factors $S' = S, S + n_1, S + 2n_1, \dots$, whereby $n_1$ denotes the blipped-CAIPI $z$-blip cycle ($S = 1$) associated to the CAIPI pattern.

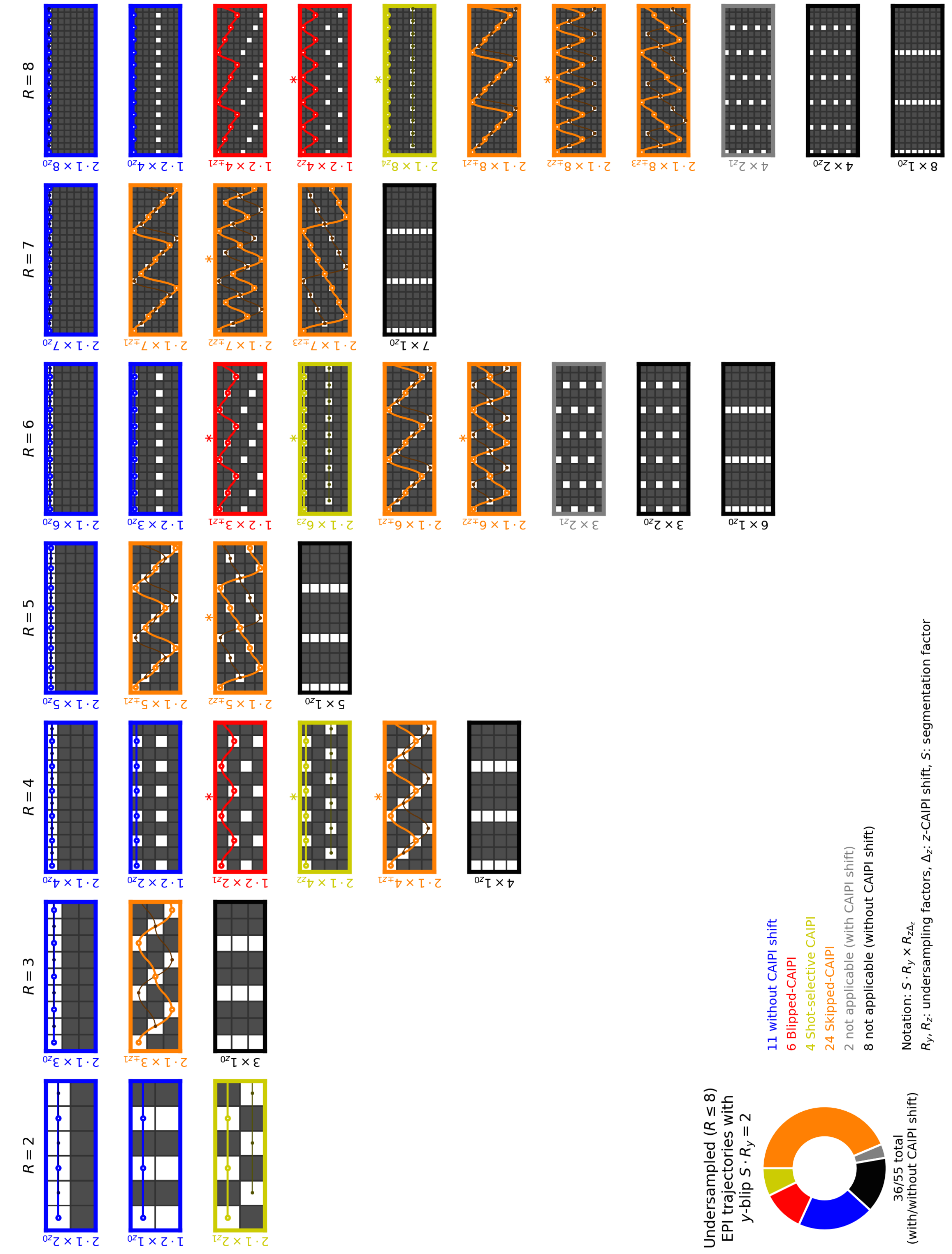



**Supporting Information Figure S3:** Applicable and not applicable EPI $R \leq 8$ undersampling with a fixed $y$-blip $S \cdot R_y = 2$. Only $\Delta_z \geq 0$ patterns are plotted, but corresponding $-\Delta_z$ patterns are counted, if applicable ($\pm$). First trajectory/subsequent trajectories indicated by thick and bright/thinner and darker curves. Sampling trajectories used for experimental time series acquisition in this work indicated by $*$ (cf. Tab. 1, Fig. 2, Supporting Information Figure S6).

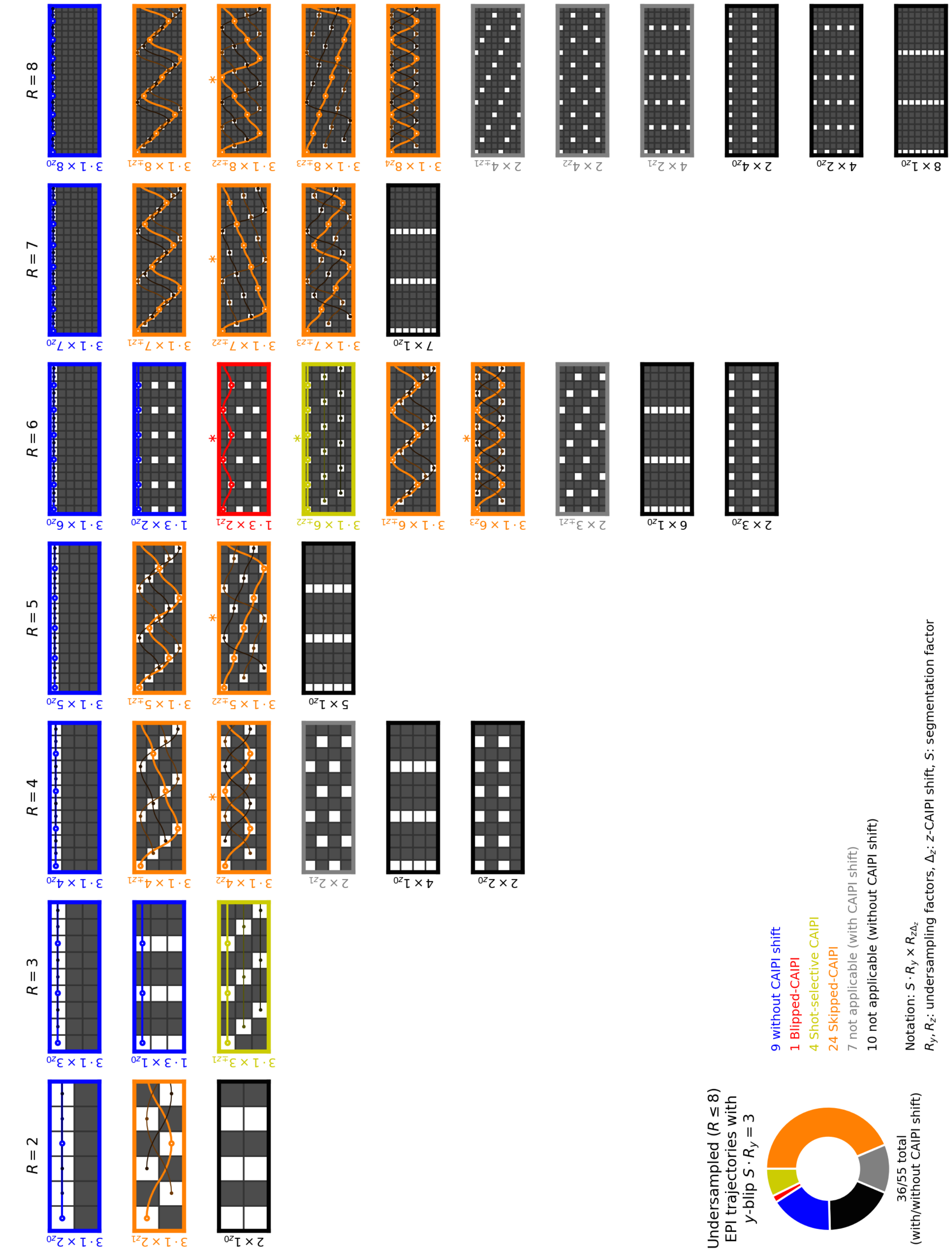


**Supporting Information Figure S4:** Same as S3 with fixed $y$-blip $S \cdot R_y = 3$.

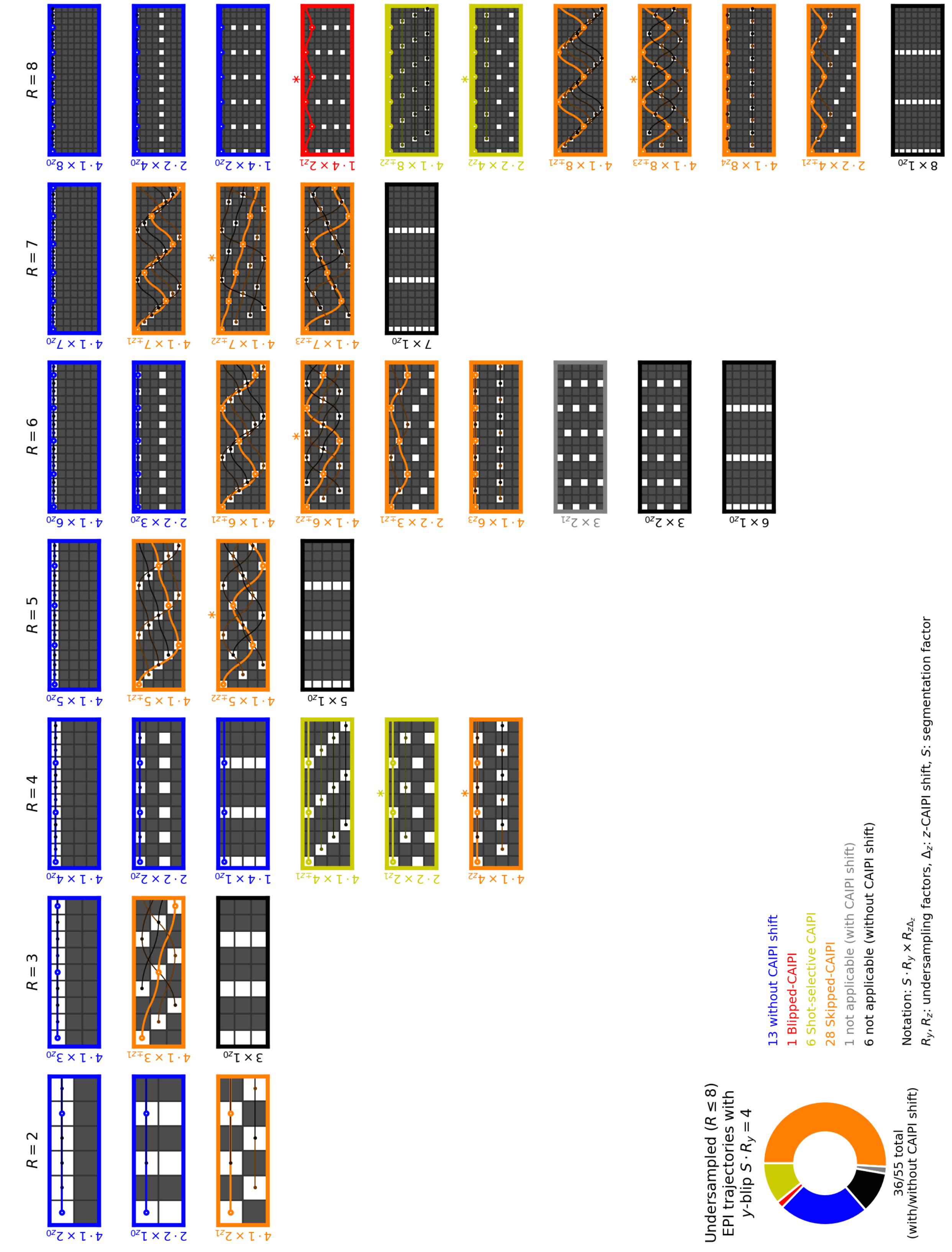


**Supporting Information Figure S5:** Same as S3 with fixed $y$-blip $S \cdot R_y = 4$. The $R = 2, 4, 6, 8$ skipped-CAIPI schemes include trajectories without $z$-blips.

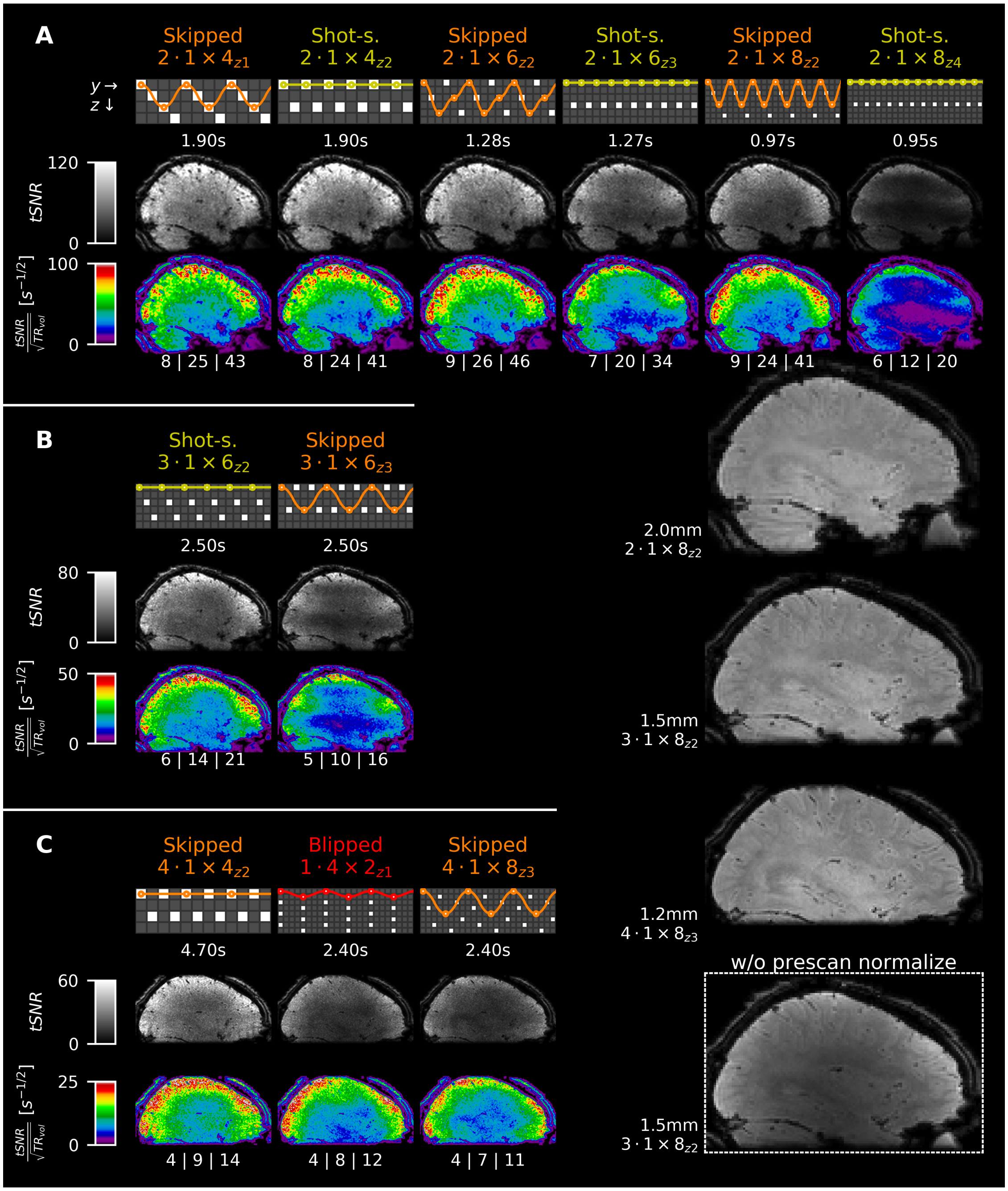


**Supporting Information Figure S6:** Temporal SNR and tSNR efficiency maps of second-to-third best time series data acquired in this work. Even though these were the best applicable blipped-CAIPI, shot-selective CAIPI and skipped-CAIPI trajectories for the respective total undersampling factors at fixed $y$-blips of 2 (A, 2mm isotropic), 3 (B, 1.5mm isotropic) and 4 (C, 1.2mm isotropic), the corresponding CAIPI patterns lead to marginally-to-substantially worse results than the optimal CAIPI patterns shown in Fig. 2. For instance, skipped-CAIPI $2 \cdot 1 \times 6_{z2}$ performed only marginally worse than the optimal blipped-CAIPI $1 \cdot 2 \times 3_{z1}$ trajectory (cf. Fig. 2). The $1 \times 6_{z3}$ CAIPI pattern performed particularly poorly, irrespective of the trajectory used (cf. A,B). However, the performance may differ using a different coil, slice coverage or slice orientation. The enlarged views on the right show the mean magnitude images of the respective $R = 8$ skipped-CAIPI acquisition. The 1.5mm example is also shown without automatic receive sensitivity correction ("prescan normalize") to indicate the receive sensitivity.

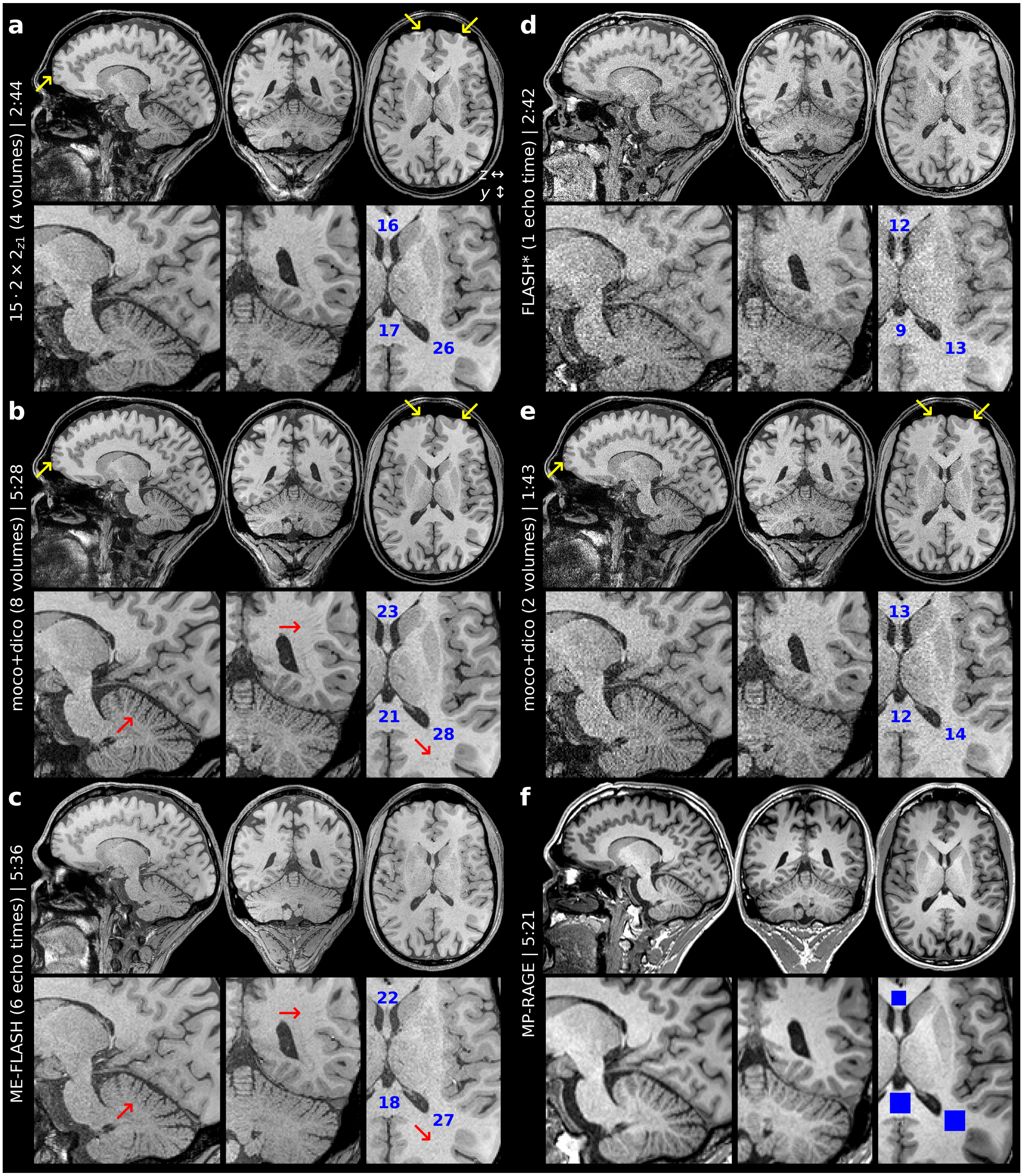


**Supporting Information Figure S7:** Orthogonal example slices of $T_1$-weighted high-resolution whole-head scans (TA stated in [min:s]) corresponding to the axial slices shown in Fig. 3 D and three additional examples (right panel). Eight magnitude-averages of the $15 \cdot 2 \times 2_{z1}$ EPI scan (b) show improved anatomical accuracy due to motion and distortion correction (moco+dico, yellow arrows) compared to four averages without corrections (a), and even improved resolution of details compared to the echo-average of a TR/FA/TA-matched ME-FLASH (c, red arrows). *A single-echo (TE=3.8ms) FLASH scan obtain with reduced TR=12ms and FA=20° but otherwise identical parameters (d) and two magnitude-averages of the $15 \cdot 2 \times 2_{z1}$ EPI following moco+dico (e) are additionally shown. Detailed views with a zoom factor of two are shown below all example slices. Blue numbers indicate rough SNR estimates from squared white matter regions indicated in the zoomed axial view of the corresponding MP-RAGE (f, 0.9mm isotropic, 240mm×240mm FOV, 192 slices, TE=2.32ms, TR=2.3s, TI=900ms, flip angle 8°, $2 \times 1$ GRAPPA, 24 integrated ACS lines, TA=5:21).

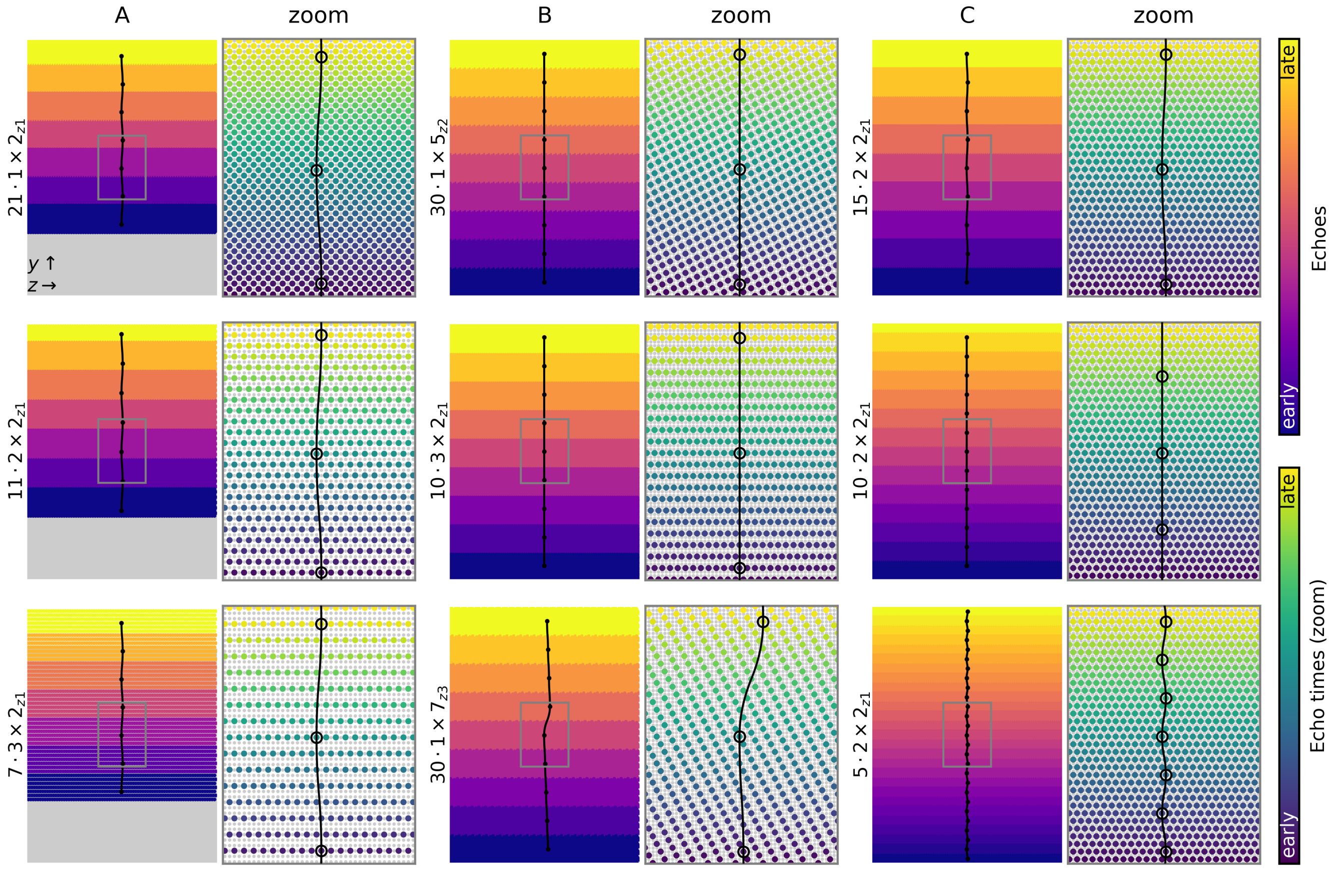


**Supporting Information Figure S8:** Visualization of the skipped-CAIPI samplings used for high-resolution $T_1$-weighted 3D-EPI scans in this work (cf. Tab. 2, Fig. 3, Supporting Information Figure S7) along with the trajectory that covers the k-space center echo (defining the image echo time, TE). Complete and zoomed k-space views use the echo number ("Echoes") and the actual echo time ("Echo times (zoom)") for color coding, respectively. Without echo time shifting (delaying the start time of each EPI trajectory by a fraction of the echo spacing according to the first primary phase encode value of the EPI trajectory, cf. Fig. 1), the echo times color-coding would resemble the step-like echoes color-coding. Image artifacts would arise, where local magnetic field inhomogeneities are strong. However, due to echo time shifting, the echo times and the MR signal magnitude and phase at each frequency-encoded echo center evolve continuously along the primary phase encode direction (1,2). As skipped-CAIPI can be understood as traditional segmented EPI with a CAIPI shift, this applies with or without CAIPI shift.
